# Machine Learning Enables Rapid Assessment of Disease Vulnerability in a Threatened Cetacean Population

**DOI:** 10.1101/2025.08.20.671308

**Authors:** Colin Murphy, Ann-Marie Jacoby, Janet Mann, Shweta Bansal, Melissa Collier

## Abstract

Cetaceans (whales and dolphins) are important ecosystem sentinels but face growing threats from major disease-related mortality events expected to intensify under climate change. Because both environmental factors (temperature, salinity) and demographics (age, sex) influence health and disease risk, understanding these relationships is essential for effective management. Direct health assessments are challenging in cetaceans, but skin lesions can indicate active infection and tooth-rake marks reflect social stressors that increase transmission risk. Yet, traditional photographic analysis of these indicators is inefficient, creating processing bottlenecks that limit timely evaluation of population health. To address this gap, we applied machine learning to rapidly assess lesions and rake marks in Tamanend’s bottlenose dolphins (Tursiops erebennus) photographed in the Chesapeake Bay, a known hotspot for disease-related die-offs. This represents the first analysis of environmental and demographic contributions to dolphin health in this region. We found significant negative relationships between lesion prevalence and both temperature and salinity for some lesion types. Adult males also showed higher rake mark coverage than adult females and calves. These patterns suggest dolphins in colder, fresher waters may face elevated disease risk, while adult males may be particularly vulnerable to behavioral stress and related health consequences. Our findings are consistent with prior studies, lending validity to our machine learning models, while also revealing novel patterns of calf and male vulnerability in this threatened population. More broadly, our approach demonstrates the potential of automated image analysis to enable timely, non-invasive health assessments across cetacean populations in an era of rapid global change.

## 1. Introduction

Wild populations of cetaceans (whales and dolphins) suffer from a variety of infectious diseases that impact their overall health and can cause mass mortalities (Simeone et al., 2015), with such events expected to intensify under climate change (Sanderson & Alexander, 2020). Many cetaceans are ecosystem sentinel species, providing early warnings about difficult-to-observe aspects of marine ecosystems (Hazen et al., 2019). Disease prevalence varies substantially among individuals and populations (Guinn et al., 2024; Hart et al., 2012; Wilson et al., 2000), driven by environmental and behavioral heterogeneity (Collier, Jacoby, et al., 2025; Collier, Urian, et al., 2025; Duignan et al., 2020; Fazioli & Mintzer, 2020; Hart et al., 2012; Wilson et al., 1999) suggesting that effective management strategies should be uniquely targeted to specific populations. However, establishing these relationships requires reliable metrics of population health, and obtaining such metrics directly (e.g., through capture–release studies) poses significant logistical and ethical challenges (Fair et al., 2014, 2017). Analyzing skin markings in photographs collected during routine field surveys offers a promising, non-invasive, alternative but the large volume of data can make processing challenging. Therefore, to rapidly enhance our understanding of factors that influence cetacean health and disease vulnerability, we developed machine learning models to quantify and characterize skin lesions and rake marks among vulnerable cetacean population(s), and analyzed their relationships with environmental and demographic factors. This approach enables us to assess disease vulnerability in real time to better inform disease surveillance and protection planning.

Skin lesions are patterns of discoloration or damage on a cetacean’s skin that are often linked to underlying diseases such as cetacean poxvirus and herpesvirus (Geraci et al., 1979; Hart et al., 2012; Maldini et al., 2010; Powell et al., 2018; Toms et al., 2020; Van Bressem et al., 1999) or environmental changes, such as major freshwater inputs or prolonged exposure (Fazioli & Mintzer, 2020; Toms et al., 2020). Evidence links lower temperature and salinity to higher lesion presence at sites along the Northwest Atlantic, including Florida, Georgia, and South Carolina (Duignan et al., 2020; Fazioli & Mintzer, 2020; Geraci et al., 1979; Hart et al., 2012; Wilson et al., 1999), suggesting that cetaceans in warmer, more saline water are less likely to suffer from skin lesions. Colder water may weaken or slow blood flow, slowing immune response (Wilson et al., 1999), while lower salinity can cause ulcers or electrolyte imbalances that compromise the integrity of cetaceans’ skin (Duignan et al., 2020; Wilson et al., 1999). These patterns thus suggest that environmental stressors may impair immune function and skin integrity in cetaceans, increasing susceptibility to infectious agents. Since skin lesions often serve as visible indicators of underlying disease, their prevalence may offer a valuable window into broader patterns of cetacean health and its relationship to their environments.

Whereas skin lesions can reflect environmental stressors or active infection, tooth rake marks can reveal how demographic traits shape health outcomes and disease vulnerability in odontocetes (toothed whales). Rake marks are patterns of parallel scarring across an individual’s skin caused by conspecific agonistic interactions and can thus be used as a metric of received aggression for individuals (Scott et al., 2005). As being the recipient of targeted aggression is often linked to immunodepression in many mammal species including primates, rodents, and pigs (Azpiroz et al., 2003), those receiving more aggression may be more vulnerable to disease and other negative health consequences (Azpiroz et al., 2003; Masud et al., 2020; Takahashi et al., 2018). Past studies have demonstrated relationships between rake marks and both sex and age. For example, in bottlenose dolphins, juveniles and adults have more rake marks than calves, and males have more rake marks than females (Lee et al., 2019; Scott et al., 2005; Serres et al., 2023) suggesting potential demographic variability in received aggression in this species. Having more rake marks could also indicate more social contacts, which is associated with greater disease risk and known to vary demographically (Collier, Jacoby, et al., 2025). Additionally, since most rake marks heal within 5-20 months (Lee et al., 2019), extreme rake mark coverage could indicate slowed healing due to nutritional stress, greater received aggression, or slower immune response (Lee et al., 2019; Serres et al., 2023). Past studies of bottlenose dolphins (*Tursiops* spp.) and killer whales (*Orcinus* spp.) also found skin lesions to be associated with rake marks, as damage to the epidermis may serve as a pathway for increased disease transmission (Gaydos et al., 2023; Toms et al., 2020). Thus, rake marks can offer an integrative indicator of social stress, immune function, and disease risk in odontocetes, and how they vary across demographics and populations.

The presence of skin lesions and rake marks on cetaceans can be assessed through photographic field data. Photo-identification studies routinely generate thousands of images per field season, creating a large volume of data that is challenging to process. Manual assessment of photographic data for identification of skin lesions and rake marks, in particular, requires extensive processing, leading to a lag between data collection and analysis. The process of identifying individual cetaceans in each image may take over 1,100 hours for a single year of data (Tyne et al., 2014), which is further compounded by the difficulty of distinguishing lesions and rake marks from each other or from other factors such as glare, splashes, or shadows. A researcher’s experience and subjectivity could further reduce interrater reliability (Toms et al., 2020). Past work addressed these issues for skin lesions by developing a machine learning tool that locates and classifies skin lesions on dolphins in photographs (Murphy et al., 2025), but no such tool yet exists for tooth rake marks.

In the current work, we apply machine learning models to analyze photographic data and identify skin lesions and rake mark presence on Tamanend’s bottlenose dolphins (*Tursiops erebennus*) inhabiting the Chesapeake Bay, U.S.. As an estuary, the Chesapeake Bay is an important and heavily utilized, though relatively underrecognized, seasonal habitat for these dolphins (Collier, Jacoby, et al., 2025; Hayes et al., 2021; Murphy et al., 2025; Rodriguez et al., 2021), and it experiences extreme fluctuations in temperature and salinity both within and between years (Baird & Ulanowicz, 1989), making it a unique system to test hypotheses relating environmental factors. Dolphins in this region have also experienced major disease-related mortality events, with morbillivirus outbreaks in 1987 and 2013 reducing some Tamanend’s bottlenose dolphin populations by 40–50% (Lipscomb et al., 1994; Waring et al., 2016). The Chesapeake Bay appeared to be a hotspot during these outbreaks (Collier, Urian, et al., 2025), and the burden was disproportionately distributed across age and sex classes (Collier, Jacoby, et al., 2025), making this population also particularly valuable for examining the demographic influences of health and susceptibility. Here, we use machine learning assessments of skin lesions and rake marks to provide the first insights into environmental and demographic drivers of dolphin health and disease susceptibility in this vulnerable region, while also demonstrating the broader potential of these models for rapid, reliable cetacean health assessments.

## 2. Methods

We used machine learning models from our past work (Murphy et al., 2025) to estimate skin lesion prevalence across photo-identification surveys of wild Tamanend’s bottlenose dolphins from the Potomac-Chesapeake Dolphin Project (PCDP) between 2015 and 2019, and developed new models to estimate rake mark coverage on individuals present in a subset of these surveys. We examined how skin lesion prevalence varies with sea surface temperature and salinity and how rake mark coverage varies across age and sex classes.

### 2.1 Photographic Data

Photo-identification surveys were conducted between the lower Potomac River (38.3616 N, 76.9971 W) and the middle Chesapeake Bay (37.8007 N, 76.2885 W) between April and October from 2015 to 2019 (Figure 1a). Researchers conducted boat-based behavioral surveys of dolphins along line transects within the study area. During behavioral surveys (hereafter referred to as surveys), observers took photographs of all present individuals and recorded their predominant behavioral activities (Karniski et al., 2015). All photographs were scored for photo quality, with zero being the lowest score (unusable quality) and three being the highest score (exceptional quality) (modified from Urian et al., 2015) (Figure S1). Each photograph was further coded for all individuals present using dolphins’ unique dorsal fin features (Hammond et al., 1990). All data were collected under NMFS General Authorization numbers 19403 and 23782.

**Figure 1.**
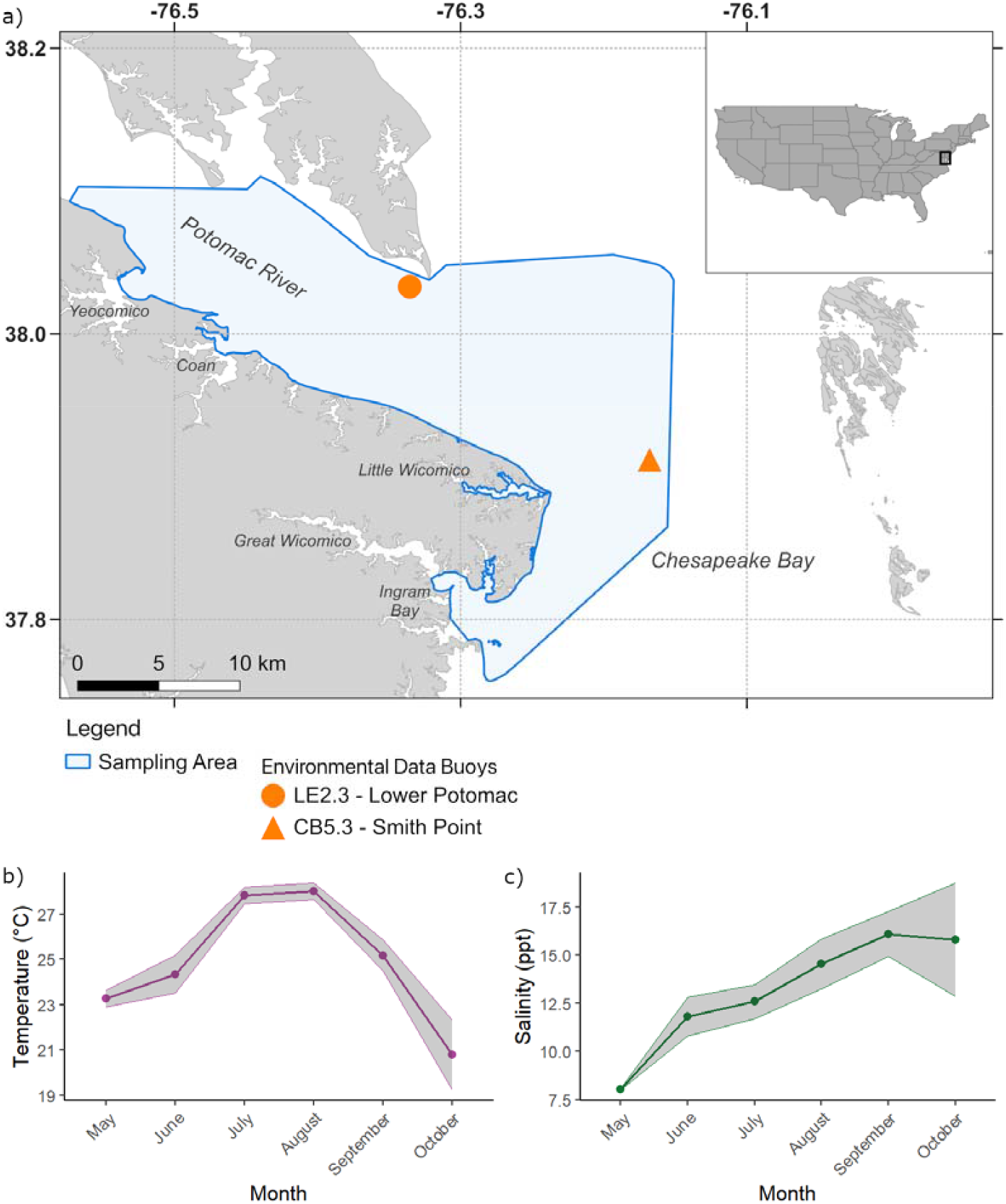
Study site and environmental data. Surveys were conducted within the Chesapeake Bay (a) between the lower Potomac River and the middle Chesapeake Bay. Environmental data from two monitoring stations within the sampling area, labeled in (a) were used to determine the average temperature (b) and salinity (c) for each survey and for each month.

### 2.2 Demographic Data

For dolphins photographed during PCDP surveys, we used a set of rules for assigning age and sex classes to each dolphin when sufficient data were available (Collier, Jacoby, et al., 2025). These rules use a combination of sighting records (e.g. presence of dependent calves across years), behavioral data (e.g. observed sexual behaviors), photographic data (e.g. confirmed genital sightings), and size observations to place individuals into one of four demographic groups: adult male, adult female, juvenile, and calf. Our rules do not allow for juveniles and calves to be confidently assigned a sex class without confirmed genital sightings.

### 2.3 Environmental Data

We obtained temperature and salinity data from the Maryland Department of Natural Resources’ Eyes on the Bay monitoring program dataset (http://www.eyesonthebay.net).

We used values from the Point Lookout monitoring station (LE2.3) (Figure 1b) in the Lower Potomac River (Maryland) and the Smith Point monitoring station (CB5.3) (Figure 1c) in the Chesapeake Bay mainstem (Virginia). These two monitoring stations are within the bounds of the PCDP study area (Figure 1a). When data were available from both Point Lookout and Smith Point on the same day, we used Point Lookout’s measurements. If both stations lacked data for a given date, we substituted values from the nearest available date at either station. For each day, we then averaged temperature and salinity across all available times and depths within the chosen dataset.

### 2.4 Estimating Lesion Prevalence

From all PCDP surveys, we filtered images to include only those with photo quality 2 (good) or 3 (excellent), and only those with a single dolphin identified in the image, as our machine learning model cannot reliably assign lesions to individuals in photographs of more than one dolphin; we did not manually crop photographs to each individual as there were greater than 50,000 images, making this logistically infeasible. As small group surveys (<5 dolphins present) were more likely to be biased towards extreme lesion prevalence estimates of 0 or 100% (Figure S3), we dropped these surveys from further analysis. Our final dataset included 15,582 images from 108 surveys (Table 1). The average number of dolphins represented in each survey was 28.42 and the full distribution can be found in Figure S4.

**Table 1.** Lesion prevalence dataset images and surveys by month and year. Data were collected between May and October, however not all months were sampled every year. Each entry indicates the number of photographs, with the number of surveys conducted in parentheses. The total by month (column) and year (row) are summed across the respective dimensions. Note while this table reflects the number of photos, we calculated lesion prevalence per survey at the level of individual dolphins, not images, to avoid pseudoreplication.

| Year | May | June | July | August | September | October | Total |
| --- | --- | --- | --- | --- | --- | --- | --- |
| 2015 | - | - | 839 (8) | - | 464 (3) | - | 1303 (11) |
| 2016 | - | 239 (2) | 162 (1) | 266 (3) | 52 (1) | - | 719 (7) |
| 2017 | - | 276 (4) | 407 (5) | 472 (4) | - | - | 1155 (13) |
| 2018 | - | 502 (8) | 480 (8) | 858 (9) | - | 221 (1) | 2061 (26) |
| 2019 | 860 (4) | 1312 (8) | 1751 (11) | 1884 (14) | 2578 (10) | 1959 (4) | 10344 (51) |
| Total | 860 (4) | 2329 (22) | 3639 (33) | 3480 (30) | 3094 (14) | 2180 (5) | 15582 (108) |

We then ran our lesion detection machine learning model (Murphy et al., 2025) to identify spot and fringe ring lesions in each photograph. Spot lesions are those that appear as simple circles, while fringe rings are lesions that appear in a concentric circle or “bullseye” pattern (Fig. 2a). Following the methodology we developed in Murphy et al., (2025) to prevent excess false positives, for dolphins captured in two or fewer photographs in a survey, we considered the dolphin lesioned if the model returned a positive lesion detection on any of its photographs. For dolphins captured in three or more photographs in a survey, we considered the dolphin to be lesioned if the model returned a positive lesion detection on more than 25% of its photographs. With each survey, we then calculated the prevalence of both spot and fringe ring lesions across all dolphins observed in that survey. Given the different and uncertain etiologies of dark versus pale spot lesions (Hart et al., 2012; Toms et al., 2020), we also calculated separate prevalence values for spot lesions results by pigmentation (pale or dark).

**Figure 2.**
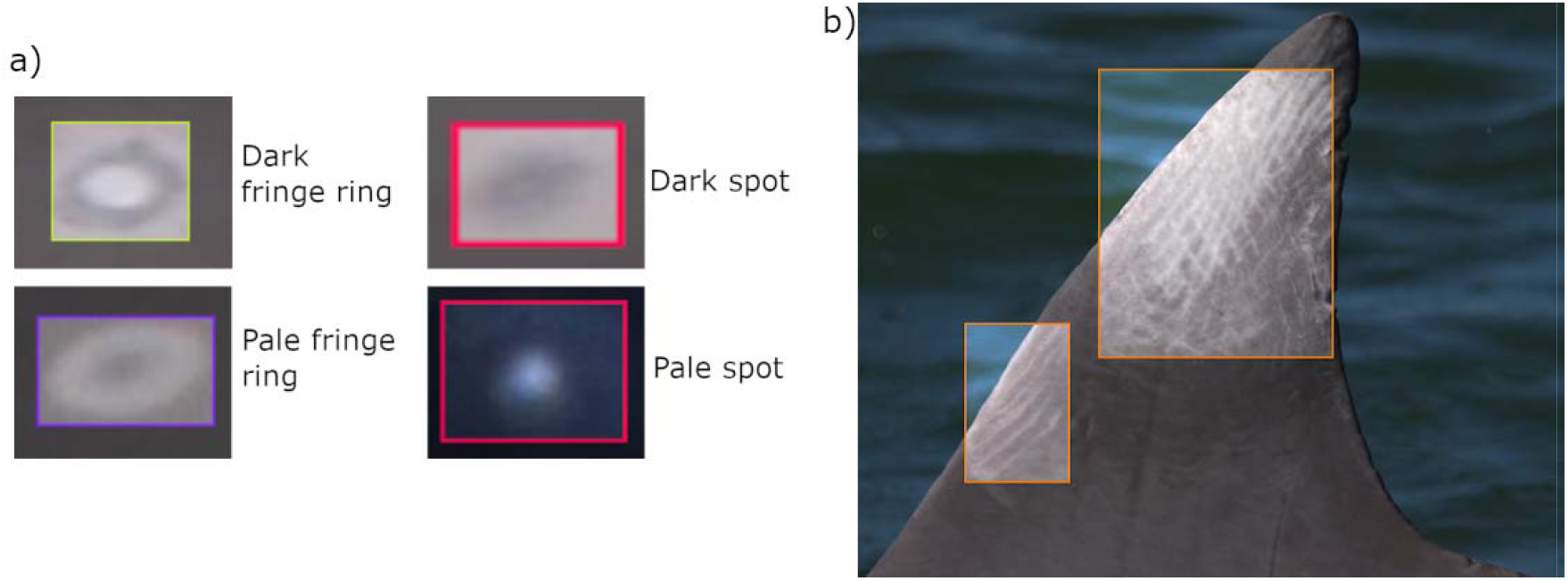
Skin markings detected by machine learning. Our machine learning models detected four different classes of lesions (a) described by their shape (fringe ring versus spot) and outermost pigmentation (pale versus dark), and rake marks (b) described by patterns of parallel scarring.

### 2.5 Estimating Rake Mark Coverage

#### 2.5.1 Developing a rake mark detection model

We created a new machine learning model capable of locating rake mark patterns from photographic data using the Roboflow Train platform (https://roboflow.com). Models constructed in Roboflow as Roboflow 3.0 object detection models are based on the site’s custom version of the YOLOv8 model developed by Ultralytics (Jocher et al., 2023). These were constructed similarly to the lesion detection models we developed previously (Murphy et al., 2025). We selected 378 images that were first cropped to the available body and dorsal fin regions using our existing body-dorsal fin detection model, an automated tool for identifying and cropping dolphin dorsal fins and body areas in photographs (Figure S5). We then developed separate models to detect rake mark coverage on the dorsal fin and body regions, respectively. This methodology reduces error from training and prediction on negative space (water or sky area) in images of dolphins (Murphy et al., 2025). The dorsal fin and body crops were randomly split into training, validation, and test datasets (consisting of 70%, 20%, and then 10% of the images, respectively) and annotated to indicate the location and size of the rake marks present in each photo (Fig. 2b).

For the dorsal fin rake mark detection models, we generated three augmented images per source image, using all augmentation techniques that were used in training the lesion detection models (Fig. S2) (Murphy et al., 2025). We used the better of two replicate models, with the highest performing model having a mean average precision (mAP) of 84.7% with the Intersect over Union (IoU) value set at 0.5, with average mAP across IoUs from 0.5 to 0.95 around 50%. This model also had a precision of 93.9% and a recall of 73.7% (see glossary for definitions of evaluation metrics). For the body rake mark detection models, we only applied mosaic augmentation as these models performed better with mosaic augmentations alone compared to models with our full set of augmentations (Fig. S2). We used the better of three replicate models, with the highest performing model having a mAP of 89.2% with the IoU value set at 0.5, with average mAP across IoUs from 0.5 to 0.95 leveling off around 73%. This model had a precision of 90.9% and a recall of 82.9%.

#### 2.5.2 Calculating Rake Mark Coverage

The rake mark detection models returned predictions of rake mark presence, position, and area for each photograph (Figure 2b). To determine the rake mark coverage on each dolphin in a photograph, we used the Python package Shapely to construct a polygon of all predicted rake bounding boxes (a representation of the visible surface area covered by rake marks), and a polygon of the predicted body and dorsal fin bounding boxes (a representation of the visible surface area of skin that rake marks could possibly cover). We then defined the extent of rake mark coverage for an individual as the area of the rake polygon divided by the area of the body and dorsal fin polygon.

#### 2.5.3 Applying the model to PCDP data

To examine rake mark coverage in PCDP dolphins, we filtered the subset of images used to determine lesion prevalence to contain only images of dolphins that have been assigned to a demographic group (Section 2.2). Due to the low sample size of confirmed juveniles present in our 2015-2019 data (n = 3), we added 12 additional juveniles that were photographed between June and July of 2021. The final dataset contained 3,958 photographs across 77 days, with 615 sightings of 108 individuals: calf (n = 8 across 22 sightings), juvenile (n = 15 across 17 sightings), adult female (n = 26 across 111 sightings), and adult male (n = 59 across 465 sightings). We determined the rake mark coverage for every sighting day for a given dolphin by averaging the coverage values across all photographs of that dolphin on that day.

### 2.6 Estimating Local Density

Because infection risk can be density-dependent, we aimed to account for potential density effects on lesion prevalence in our analysis. Given the fission-fusion social structure of bottlenose dolphins (Connor et al., 2000), we first aimed to identify an appropriate metric of “localized density”, defined as the close (<10m) contacts experienced on average by an individual in a survey. We compared survey size (total number of dolphins photographed during a survey) with the average number of dorsal fins per photograph in a survey (as assessed using our dorsal fin detection model (Figure S5), using data on close contacts from 67 individual focal follows that were associated with 59 surveys as an independent reference for localized survey density (see supplementary information for more details). Both measures correlated significantly with localized density, but average fins per image showed a stronger relationship, so we used it as our measure of localized density in our analysis (Figure S6).

### 2.7 Statistical Analyses

To assess the relationship between skin lesion prevalence, temperature, and salinity, we constructed a generalized linear mixed model (GLMM) where a survey’s lesion prevalence was the response variable, the temperature and salinity for each survey were included as predictor variables, and the date of the survey was added as a random effect. We controlled for the potential effects of density-dependence on lesion prevalence, using the average number of dorsal fins per image in each survey as a fixed effect (section 2.6). We constructed three total models to assess the impact of temperature and salinity on 1) dark spot prevalence, 2) pale spot prevalence, and 3) fringe ring prevalence. The models were run in R using the lme4 library.

To examine the relationship between demographics and tooth rake coverage, we used a GLMM where an individual’s daily rake mark coverage was the response variable, their demographic class (adult male, adult female, juvenile, calf) on that day was a categorical predictor variable, the year was an additional fixed predictor, and the dolphin’s identity was a random effect. The models were run in R using the lme4 library.

## 3. Results

### 3.1 Environmental factors predict fringe ring-class lesion prevalence and pigmentation-specific spot-class lesion prevalence

The mean lesion prevalence observed across all surveys was 72.3% (n = 108, 95% CI = [69.2%, 75.4%]). In other words, on average, 72.3% of dolphins in a survey presented one or more skin lesions. We found significant associations between both temperature (p=4.85e-04) (Fig. 3a, Table S2) and salinity (p=7.66e-05) (Fig. 3d, Table S2) and fringe class-specific lesion prevalence. When we investigated trends separately for pale spots and dark spots, we found no significant association between temperature and either spot class lesion type (p=0.751, Fig. 3b for pale spots; p=0.101, Fig. 3e for dark spots; Table S2 for both). However, we did find a significant positive association between pale spot prevalence and salinity (p=4.51e-04, Fig. 3c, Table S2), and a significant negative association between dark spot prevalence and salinity (p=1.32e-03, Fig. 3f, Table S2). No evidence was found for density-dependence for any lesion class (Table S2).

**Figure 3.**
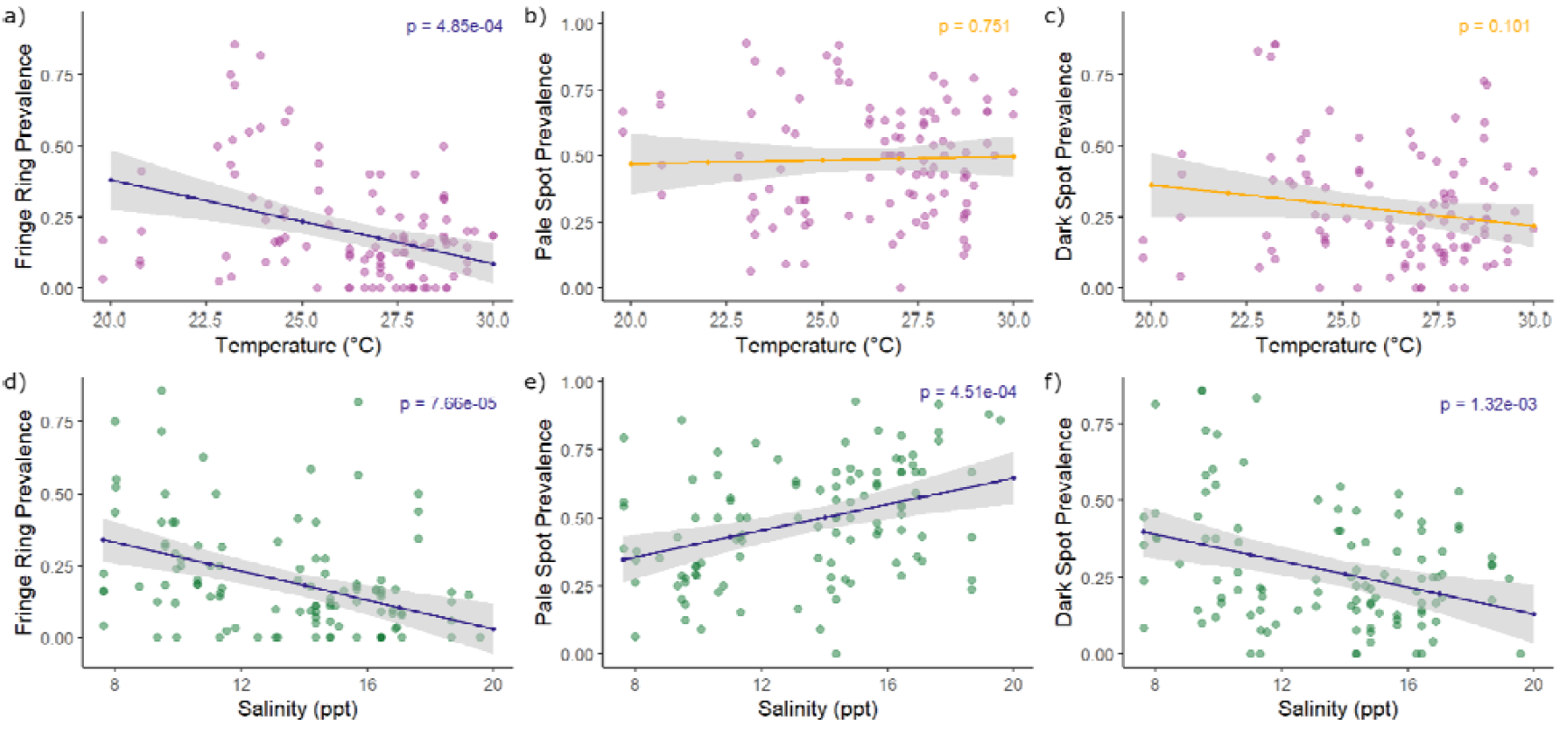
Environmental factors show significant association with lesion prevalence across most lesion classes. The effect of temperature on a) fringe ring lesion prevalence, b) pale spot lesion prevalence, and c) dark spot lesion prevalence, as well as the effect of salinity on d) fringe ring lesion prevalence, e) pale spot lesion prevalence, and f) dark spot lesion prevalence. Dots represent the individual survey observations and lines represent glmm predictions with the gray area representing the 95% confidence interval. Blue lines represent a significant association, while orange lines represent an insignificant association. Temperature was significantly associated with fringe ring prevalence, and salinity was significantly associated with the prevalence of all lesions (n= 108 surveys).

### 3.2 Adult males have higher rake mark coverage than other demographic classes

The average rake mark coverage on a dolphin’s visible body surface for a given date was 4.34% (n = 615, 95% CI = [3.91%, 4.78%]). We found that the adult males had significantly higher rake mark coverage than adult females (p=9.23e-04) and calves (p=6.11e-03) (Fig. 4, Table S3). We found no significant difference in rake mark coverage between adult female dolphins and calves (p=0.368) or juveniles (p=6.80e-02), but we found that juveniles had significantly higher rake mark coverage than calves (p=3.16e-02) (Fig. 4, Table S3).

**Figure 4.**
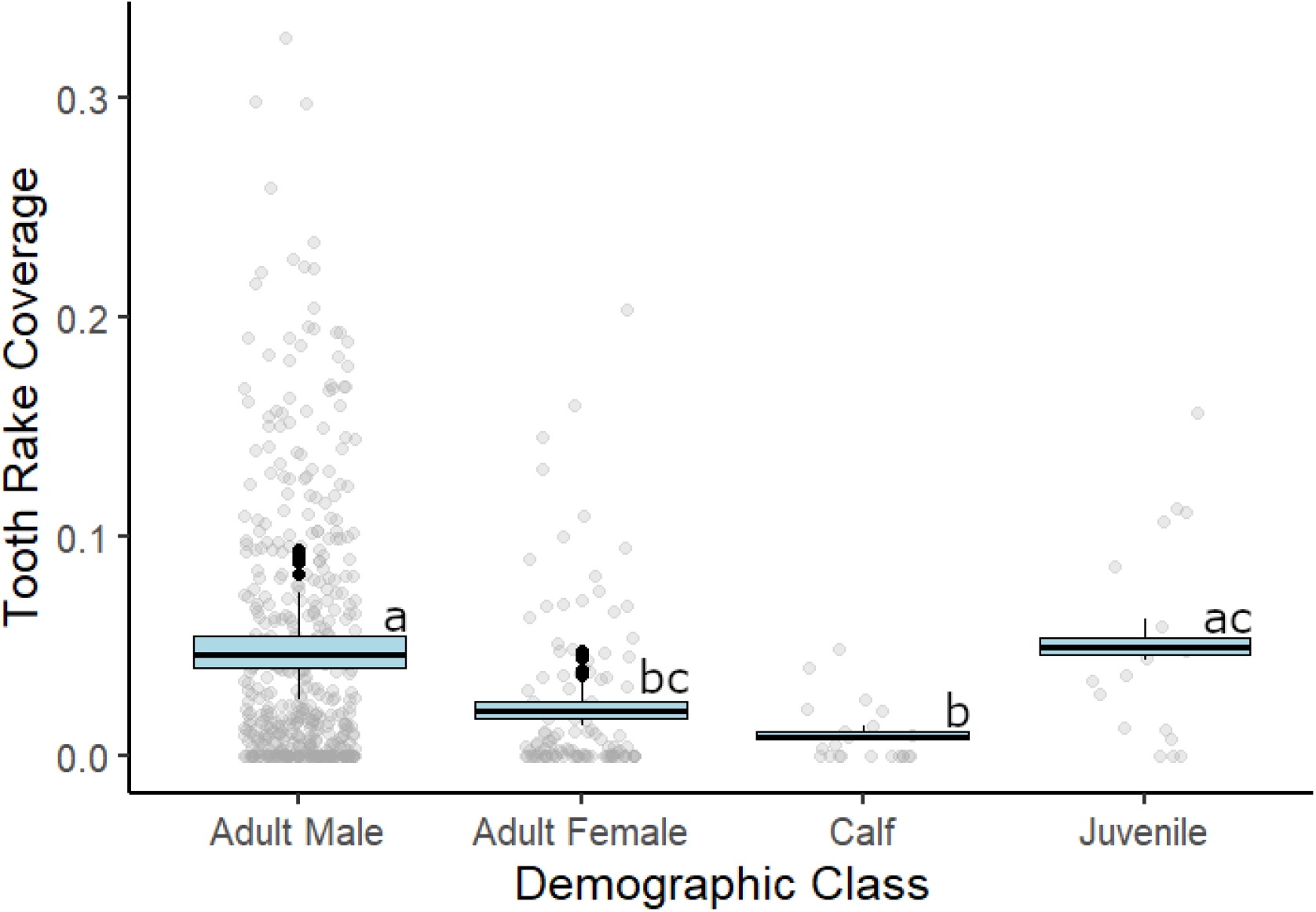
Adult males show significantly higher rake mark coverage than adult females and calves. Gray dots represent raw data as a coverage estimate for individuals for each day sighted, boxplots represent model estimates for each demographic group, and letters indicate significant differences. Adult males had significantly higher rake mark coverage than adult females and calves, but not juveniles.

## 4. Discussion

We applied a new machine learning technique to efficiently assess the effects of environmental and demographic influences on skin lesion and tooth rake mark prevalence among vulnerable dolphins in an understudied habitat (Collier, Jacoby, et al., 2025; Murphy et al., 2025). We found that fresher, lower salinity water was associated with higher prevalence of fringe ring lesions, while results varied for temperature. We also found that adult males have significantly higher rake mark coverage than adult females and calves; adult females and calves do not differ in rake mark coverage; juveniles were not different from adult males. By combining our machine learning tools with environmental and demographic data, we provide new insights into health trends for Chesapeake Bay dolphins, and highlight how skin lesion and rake mark patterns can be used to provide rapid assessments of spatial and social variation in health and disease vulnerability among cetaceans.

We observed varying environmental trends of prevalence across different lesion classes. The etiologies of these lesion classes vary; spot lesions are not typically of infectious origins, while to fringe ring lesions have been closely linked to cetacean poxvirus and herpesvirus (Hart et al., 2012; Maldini et al., 2010; Toms et al., 2020); pale spots have been occasionally tied to cetacean herpesvirus (Hart et al., 2012; Toms et al., 2020) but the etiology of dark spots is unknown (Toms et al., 2020). Temperature was not associated with pale and dark spot lesions, but had a significant negative relationship with fringe ring lesions. Given the close relationship between fringe ring lesions and disease, our results suggest heightened disease vulnerability in colder waters for dolphins in this region, consistent with past studies (Hart et al., 2012; Wilson et al., 1999). Both marine mammals and their prey species have begun to exhibit poleward shifts in their habitat ranges (Poloczanska et al., 2016); given that the Chesapeake Bay is part of their northern most range (Hayes et al., 2021), Tamanend’s bottlenose dolphins may stay in colder waters for longer than they have previously. However, a recent study showed increased disease mortality at higher sea surface temperatures (Williams et al., 2025), indicating that trends in lesion prevalence may be site-specific or contrary to trends in actual mortality. Our data were focused primarily in the summer months such that colder water surveys were underrepresented, and as such additional data could provide better insights into the trends observed in this research. Spot lesions also have an extremely high prevalence overall among dolphins in the Chesapeake Bay compared to the fringe ring-class lesions (Murphy et al., 2025), indicating that a higher sample size may be required to detect temperature trends with spot lesions.

We found that the prevalence of both fringe ring and dark spot lesions trended negatively with salinity. This is consistent with past studies describing links between lowered salinity and increased skin lesion prevalence (Duignan et al., 2020; Fazioli & Mintzer, 2020; Hart et al., 2012; Wilson et al., 1999). This is especially important to consider given climate-induced increases in the intensity of storms (Kossin et al., 2020), which may lead to further flooding and freshwater intrusion in marine systems. However, we also found that pale spot lesions demonstrated a positive association with salinity, such that surveys conducted when waters were more saline had a higher average prevalence of dolphins with pale spot lesions than surveys conducted in fresher waters. While this relationship was unexpected based on past research, it is important to note that both temperature and salinity may serve as proxies for other seasonal changes, such as migratory patterns or social behaviors that fluctuate predictably with the seasons, and additional research should be conducted to determine the exact relationship between these variables. For example, multiple populations of migratory Tamanend’s bottlenose dolphins utilize the Chesapeake Bay at different times of year (Collier, Urian, et al., 2025; Hayes et al., 2021; Waring et al., 2016), thus generating a better understanding of which populations are present each month and where else along the coastline they visit may allow us to better understand environmental trends in lesion prevalence and disease vulnerability (Taylor et al., 2021). In addition, the Chesapeake Bay is a brackish system with salinity levels already well below those of oceanic waters (Baird & Ulanowicz, 1989), underscoring that dolphins in this region may already experience heightened physiological stress. Together, our findings highlight the need to disentangle the direct effects of temperature and salinity from seasonal drivers to better predict how changing environmental conditions will shape disease vulnerability in this population.

We found that adult males in the Chesapeake Bay have significantly higher rake mark coverage when compared with adult females, and both adult males and juveniles have significantly higher rake mark coverage than calves. These findings align with *Tursiops* studies elsewhere (Lee et al., 2019; Scott et al., 2005; Serres et al., 2023), which found that males had both higher prevalence and intensity of rake marks compared to females. This is likely due to high rates of mating competition among males in this gregarious genus, where access to females is vigorously contested (Connor & Whitehead, 2005; Cunningham & Birkhead, 1998; Scott et al., 2005). Interestingly, we also found that calves do not have significantly lower rake mark coverage than adult females, in contrast to past work (Lee et al., 2019; Scott et al., 2005). This could be explained by unique trends in behavior in large populations of dolphins in the northern Atlantic Ocean, where calf-directed aggression may be higher compared to other populations where such rake mark studies have been carried out (McEntee et al., 2023). In another study conducted in the Potomac-Chesapeake region, 18% of very young calves (< 3 months) had tooth rake marks, significantly more than another population in Shark Bay, Australia (Long et al. in prep). Such aggression can have immunosuppressive effects (Azpiroz et al., 2003), and high tooth rake coverage may also reflect slower healing (Lee et al., 2019; Serres et al., 2023), higher rates of contact, or provide additional pathways for pathogen entry (Gaydos et al., 2023; Toms et al., 2020). Therefore, our results suggest that males may be especially vulnerable to disease outbreaks, and that calves in the Chesapeake Bay region may face heightened disease risk compared to calves elsewhere. Given that rake mark prevalence and extent is known to vary across populations due to differences in sociality, habitat, and season (e.g. Serres et al., 2023), these findings highlight the importance of considering both ecological context and social behavior when interpreting tooth rake marks as indicators of health and risk in wild dolphin populations.

Effective monitoring of cetacean health is not a one-size-fits-all approach. Typically, it requires targeted monitoring unique to populations to anticipate periods and individuals with increased disease risk. Our findings highlight key environmental and demographic factors that may influence disease vulnerability of Tamanend’s dolphins in the Chesapeake Bay. We suggest that surveillance efforts in this region could be tailored to focus on colder seasons with higher rainfall when individuals may be more vulnerable to disease. While disease intervention strategies remain logistically challenging for most cetacean species, research has shown that human disturbances can worsen disease outcomes in marine mammals (Collier et al., 2022), and thus managing disturbance levels during these vulnerable periods could help mitigate some infection. Additionally monitoring freshwater inputs into the Chesapeake Bay from runoff, rainfall, and major storm events will continue to be important, especially as climate change is expected to increase the intensity of major precipitation events (Kossin et al., 2020). Our results found that low salinity is also associated with a greater incidence of lesions indicative of poxvirus, a skin disease that is known to weaken immune systems (Van Bressem et al., 2022) thus increasing vulnerability to other more deadly disease such as cetacean morbillivirus, responsible for major die-offs of cetaceans worldwide (Lipscomb et al., 1994; Van Bressem et al., 2014; Waring et al., 2016). Past work has found that adult males are particularly vulnerable to outbreaks of morbillivirus given their high rates of assortative contact (Collier, Jacoby, et al., 2025), and our work can provide additional explanations for these male biased infection rates which are also commonly observed in a variety of other species (Collier, Jacoby, et al., 2025; Ferrari et al., 2004; Godfrey, 2013; Silk et al., 2018). Variations in infection rates and health across demographic groups can also influence population recovery following major die-offs. For instance, our findings suggest that the heightened vulnerability of calves in the Chesapeake Bay could mean that these populations require more time to rebound from morbillivirus epidemics that have previously halved their numbers (Lipscomb et al., 1994; Waring et al., 2016). Our automated approach allows managers to assess how health and vulnerability to disease vary environmentally and demographically in real time, allowing for immediate consideration of these factors in surveillance, management, population, and climate assessments.

While many of our findings align with patterns previously reported in the literature, this study provides the first health and disease assessment for bottlenose dolphins in the Chesapeake Bay, a known hotspot for mass dolphin mortality events and a region with substantial environmental variability. Equally important, we demonstrate that machine learning can rapidly and non-invasively process large photographic datasets to generate these assessments. The consistency of our results with existing studies supports the potential validity of this approach, highlighting its promise for efficiently assessing disease vulnerability across cetacean populations and informing timely health assessments and conservation responses.

### Glossary: Machine learning terminology

Definitions of relevant machine learning terminology used in assessing machine learning model performance.

| Term | Definition |
| --- | --- |
| True Positive | The model correctly predicted a lesion where one was present. |
| True Negative | The model correctly failed to predict a lesion where one was absent. |
| False Positive (Type I Error) | The model incorrectly predicted a lesion where one was absent. |
| False Negative (Type II Error) | The model incorrectly failed to predict a lesion where one was present. |
| Precision | The number of true positives divided by the sum of true positives and false positives ($\frac{-}{+}$); when the model predicted a lesion, how often was it correct? |
| Recall | The number of true positives divided by the sum of the true positives and the false negatives ($\frac{-}{+}$); when a lesion was present, how often did the model predict a lesion? |
| Mean Average Precision (mAP) | The average of the area underneath a precision-recall curve (a plot that visualizes the tradeoff between precision and recall) across all lesion classes for a model. |
| Intersect over Union (IoU) | The area of overlap between a predicted bounding box and a ground truth bounding box divided by the total area of the two shapes combined ($\frac{-}{+}$). Used to assess how close a prediction is to the ground truth. |

## Acknowledgements and Funding

We thank Roboflow for providing access to their platform for model creation and assessment. We would also like to thank Eric Patterson and Megan Wallen for their assistance with data collection, and Milan Dolezal, Amelia Smith, and Katherine Dammer for their assistance with lesion coding on our dataset. All data from the PCDP was collected under NMFS Permit nos 19403 and 23782. This work was supported by the Morris Animal Foundation Award #D24ZO-425, and funding from Georgetown University. Data collection for this work by the PCDP was also supported by the Potomac River Keepers, the Rogers Family Foundation, the Campbell Foundation, the Scheidel Foundation, Georgetown University Earth Commons, Green-Rosenblum Family Foundation, the Wildlife Conservation Society, the National Geographic Society Grant WW-022ER-17, Waldorf Toyota, and individual donors.

