## Supplementary Information for "Machine Learning Enables Rapid Assessment of Disease Vulnerability in a Threatened Cetacean Population"

**Figure S1.** **Image quality scoring for photographs in the PCDP.** Grading criteria (focus, visibility, contrast, and angle) from Urian et al., 2015. Table from Dolezal et al., 2023. While the figure is specific to analyses of the dorsal fin, we used these criteria for all visible regions of the dolphin in each photograph.

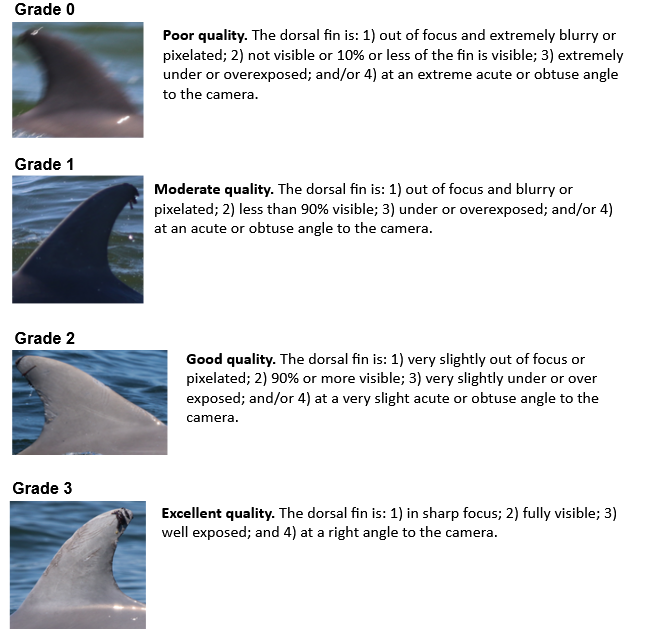

**Figure S2. Augmentation styles used in training rake mark detection machine learning models.** Models optimized to detect rake marks in images of a dolphin’s dorsal fin used all six augmentation types pictured below (0-20% crop, -15^o^ to 15^o^ rotation, -5^o^ to 5^o^ shear, 0 to 0.5 pixel blur, 0 to ~1% noise, and mosaic augmentations). Models optimized to detect rake marks in larger images of a dolphin’s body used only the mosaic augmentations. Figure from Murphy et al. (2025).

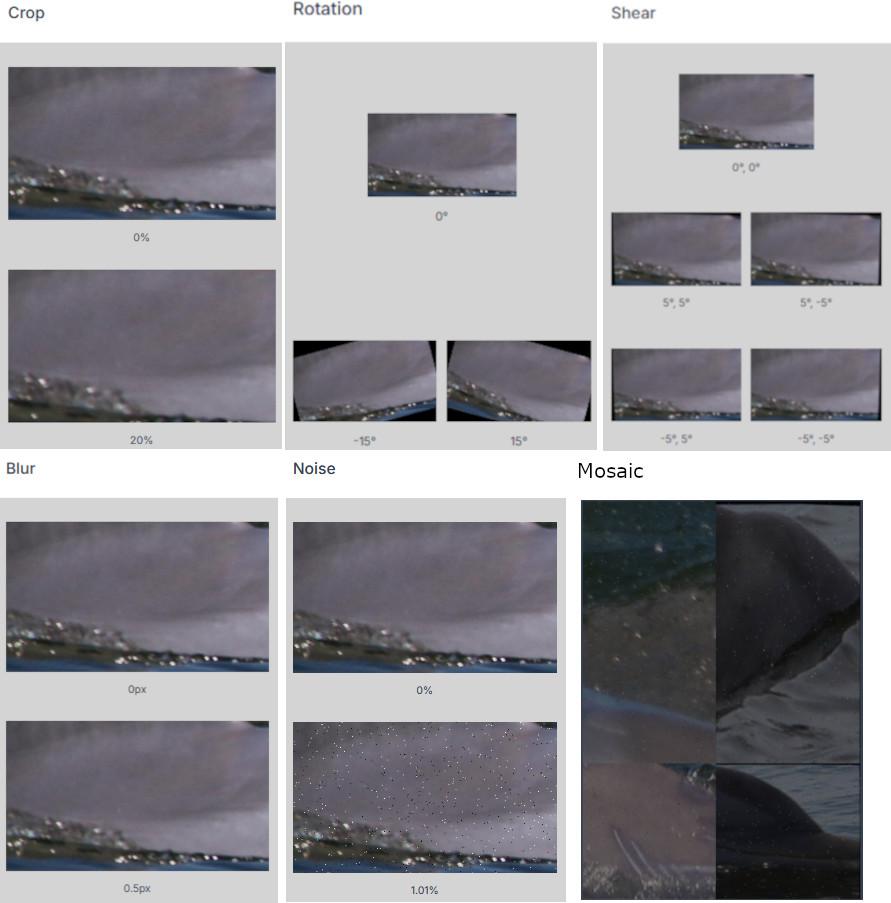

**Figure S3. Smaller surveys display bias in prevalence.** The orange dots represent individual surveys, where the number of individual dolphins observed in the survey is plotted against the lesion prevalence predicted by our models. The red line represents a simple linear model fitted to the data, and the gray shaded area is a 95% confidence interval around the linear model. We found that surveys with fewer dolphins observed (<5 dolphins) were more likely to be biased towards prevalence estimates of 0% or 100%, and as such these small surveys were dropped from further analysis.

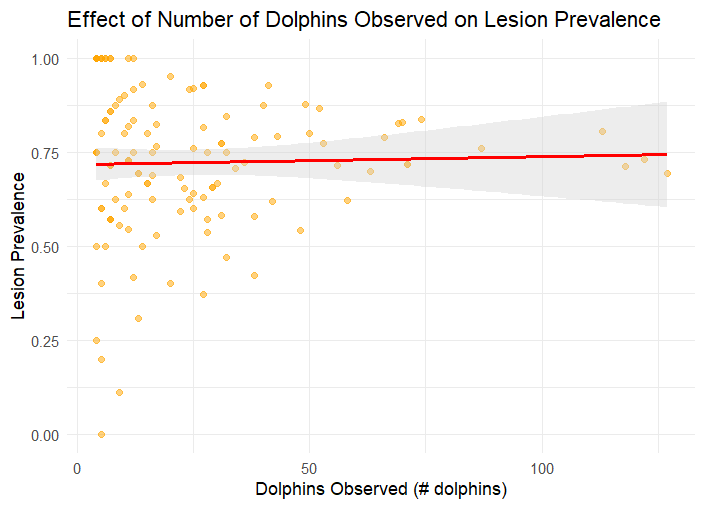

**Figure S4. Dolphins represented per survey by month.** The average number of dolphins represented in each survey used for analysis was 28.42 after dropping surveys with <5 dolphins observed.
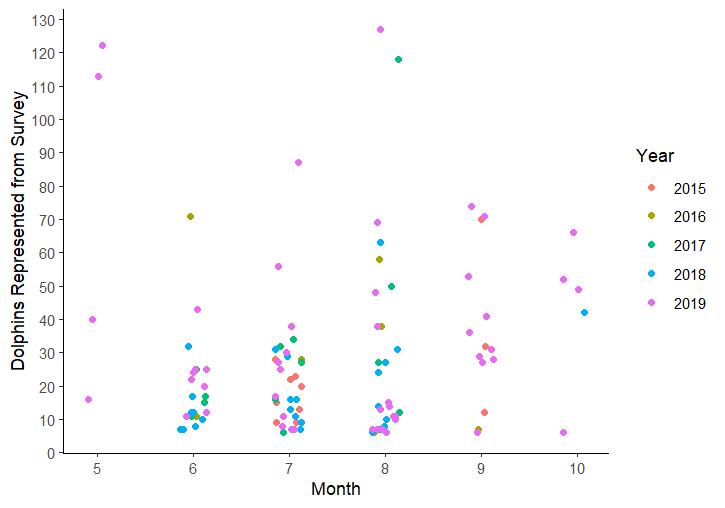

**Figure S5. Results from body/dorsal fin detection models.** As a critical element in our detection workflow, field images are initially passed to a model that automatically detects dolphins present within an image, and then passes the bounding boxes as images to the lesion or rake mark detection models. We also used this model to count the number of dorsal fins across all images for each survey to produce our proxy for density below.

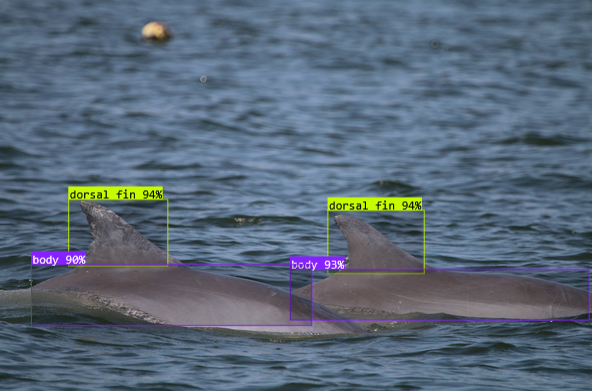

**Table S1. Breakdown of images used for rake mark coverage analysis.** Our data ranges from May through October, though not every year included such broad sampling. Each entry indicates the number of photographs, with the number of surveys in parentheses. The row labeled total indicates the sum of observations for the same month across different years, while the column labeled total indicates the sum of observations from a single year across all months included.

| **Year** | **May** | **June** | **July** | **August** | **September** | **October** | **Total** |
| --- | --- | --- | --- | --- | --- | --- | --- |
| **2015** | - | - | 175 | - | 195 | - | 370 |
| **2016** | - | - | - | - | - | - | - |
| **2017** | - | 86 | 40 | 137 | - | - | 236 |
| **2018** | - | 158 | 68 | 302 | - | 13 | 552 |
| **2019** | 53 | 503 | 391 | 563 | 830 | 414 | 2772 |
| **2020** | - | - | - | - | - | - | - |
| **2021** | - | 10 | 20 | - | - | - | 30 |
| **Total** | 53 | 757 | 694 | 1002 | 1025 | 427 | 3958 |

**Table S2. Environmental factors predicting lesion prevalence.** This table represents the output from the GLMMs constructed to predict the prevalence of fringe ring, pale spot, and dark spot skin lesions. The estimated water temperature, salinity, and average density for each survey were included as fixed effects, and the date of each survey was included as a random effect. The table was generated by the tab_model method in the sjPlot package for R (Lüdecke, 2024) using Satterthwaite's method.

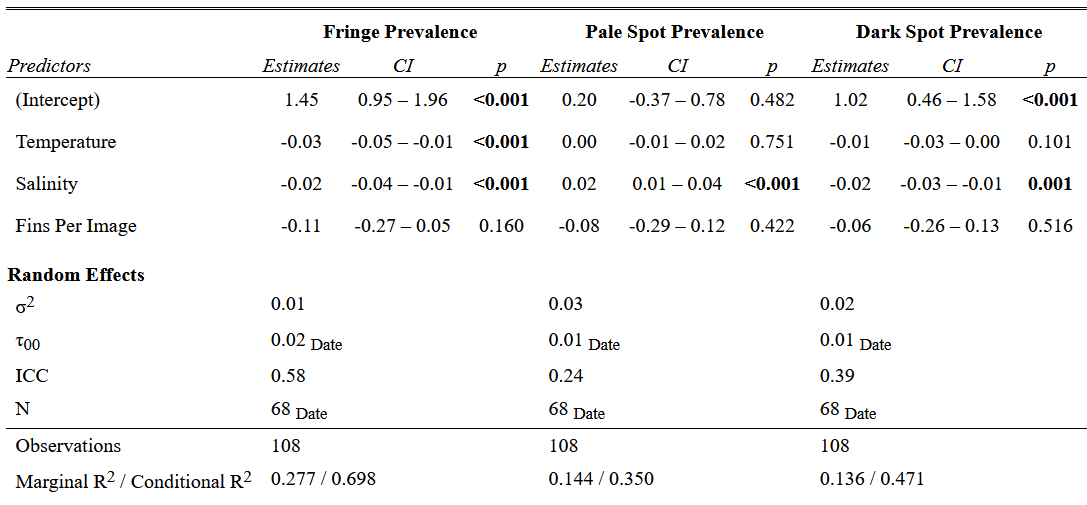

**Table S3. Demographic factors predicting extent of rake mark coverage.** This table represents the output from the GLMMs constructed to predict the extent of rake mark coverage for a sighting of a dolphin. The dolphin’s demographic class (adult male, adult female, juvenile, or calf) and the year the sighting took place were included as fixed effects, and the identity of the dolphin sighted was included as a random effect. The table was generated by the tab_model method in the sjPlot package for R (Lüdecke, 2024) using Satterthwaite's method.

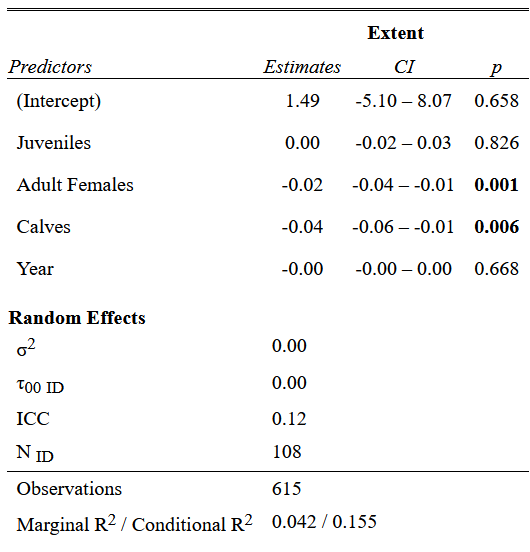

**Evaluating Localized Density Metrics**

Because infection risk can be density-dependent, we sought to control for potential density effects on lesion prevalence in our models. Given the fission-fusion social structure of bottlenose dolphins, we aimed to identify an appropriate metric of “localized density”, defined as the close (<10m) contacts experienced on average by an individual in a survey. While the survey size is recorded for each PCDP survey, this metric may overestimate localized density for individuals; for example, a survey might document 50 dolphins in the broader area, but this does not necessarily reflect the actual subgroup size or contact rates experienced by each dolphin. We therefore compared two potential metrics of localized dolphin density: (1) the recorded survey size (i.e., the total number of dolphins observed in a survey) and (2) the average number of dorsal fins per photograph within a survey, as determined by our dorsal fin detection model.

To obtain a ground-truth measure of localized density, we used data from 67 focal follows associated with 59 surveys conducted between 2021–2022. For each focal individual, we recorded their average number of close associates (within 10 m) observed throughout the follow (“follow size”). Because each focal follow was associated with a corresponding survey, follow size provides an independent measure of the true localized density within the associated survey.

We then assessed correlations between follow size and each of the two metrics: survey size and average number of dorsal fins per photograph. The left panel of Figure SX shows the correlation between survey size and follow size (r = 0.33, p = 0.006), while the right panel shows the correlation between average fins per image and follow size (r = 0.56, p<0.0001). While both metrics show significant positive correlations with follow size, the average number of fins per image demonstrated a stronger correlation, suggesting that it is a more reliable indicator of localized density than total survey size. We thus use this metric in our generalized linear models for assessing lesion prevalence to control for potential effects of density dependence.

We note that for both comparisons, outliers occurred more often at larger follow sizes, particularly when survey size and average fins per image were also high. This pattern suggests that larger subgroups may be more accurately reflected by both survey size and average fins per image.

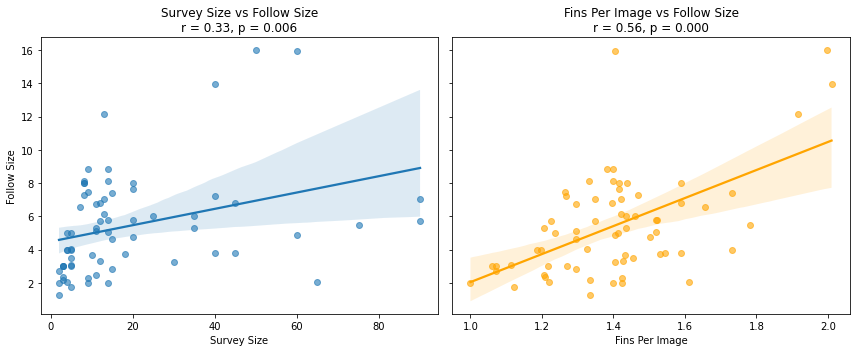

**Figure S6. The correlation between follow size and measures of localized density.** Assuming follow size to be a ground truth metric of localized density, we assess the correlation between follow size and survey sizes (left) and the average number of dorsal fins per image (right) for each survey associated with that follow. We find that the average number of dorsal fins per photo has a stronger correlation with follow size, and is thus a better metric of localized density.
